# Chemogenetic Inhibition of the Cortical Amygdala Reduces Alcohol Intake and Restores Thalamic Connectivity in Dependent Female Mice

**DOI:** 10.64898/2026.05.07.723549

**Authors:** Tiange Xiao, Xi Cheng, Jingliang Zhang, Yueyi Chen, Zhefu Que, Xiaoling Chen, Danielle McAuliffe, Alyssa Boisvert, Yang Yang, Alexander Chubykin, Adam Kimbrough

## Abstract

**Background:** Alcohol use disorder is a chronic relapsing condition characterized by excessive drinking and withdrawal symptoms. Alcohol dependence disrupts function across multiple brain regions, and recent evidence implicates the cortical amygdala (CoA) as a critical node in alcohol-related circuits. However, how CoA activity influences alcohol intake and brain-wide network function during withdrawal remains unclear.

**Methods:** Alcohol dependence was induced using chronic intermittent ethanol vapor (CIE). In one cohort, electrophysiological activity of CoA neurons was assessed during withdrawal. In a second cohort, mice underwent CIE paired with two-bottle choice drinking, and inhibitory DREADDs (hM4Di) were used to suppress CoA activity during drinking and withdrawal while behavioral outcomes were measured. Brains were then collected for Fos immunolabeling and iDISCO+ based whole-brain activity mapping to determine how CoA inhibition during withdrawal altered network organization.

**Results:** Repeated CIE increased alcohol sensitivity in CoA neurons during withdrawal. Chemogenetic inhibition of the CoA reduced alcohol intake in dependent mice without affecting withdrawal-related behaviors. Whole-brain Fos mapping showed that CoA inhibition reduced activity within the CoA while enhancing functional connectivity across multiple brain regions, particularly in the isocortex, thalamus, and anterior hypothalamic nucleus. During withdrawal without CoA inhibition, thalamic regions exhibited negative connectivity, consistent with disrupted network function; CoA inhibition reversed this pattern, producing strongly positive thalamic and medial prefrontal cortex connectivity.

**Conclusions:** These findings demonstrate that alcohol dependence alters CoA sensitivity, alcohol dependence-induced drinking and brain-wide network organization during withdrawal. The CoA appears to selectively regulate withdrawal-associated alcohol drinking, and its inhibition may reduce intake by restoring thalamic and cortical connectivity.

**Highlights:**

1. This study identifies the cortical amygdala as a previously underexplored brain region involved in alcohol-related behaviors.
2. By integrating chemogenetic inhibition with brain-wide network analysis, the study reveals candidate circuit connections through which the CoA may regulate alcohol dependence-related brain activity.
3. This study establishes the CoA as a potential driver of excessive alcohol drinking and alcohol-related network dysfunction.

## Introduction

Alcohol use disorder (AUD) is a chronic relapsing condition characterized by compulsive alcohol consumption, impaired control over intake, and the emergence of negative affective and somatic states during alcohol withdrawal (Koob et al., 2021; Koob, 2020; Koob, 2024; Koob and Le Moal, 2001). AUD represents a substantial public health burden, affecting more than 28.9 million individuals in the United States and contributing to over 140,000 alcohol-related deaths annually (NSDUH, 2024). Alcohol consumption associated with AUD ranges from episodic binge drinking to sustained heavy use, with escalation increasingly driven by withdrawal-associated negative affect and stress-related processes Alcohol (Koob, 2024; Koob and Volkow, 2016a). These features of AUD are modeled preclinically using distinct rodent behavioral paradigms that produce alcohol dependent phenotypes and withdrawal signs (Becker, 2000; Vierkant et al., 2023)

Multiple cortical and limbic regions contribute to alcohol-related behaviors (Koob and Volkow, 2016a). Increasing evidence indicates that alcohol dependence disrupts coordinated activity across large-scale brain networks rather than isolated brain regions (Kimbrough et al., 2020b; Maleki et al., 2022; Roland et al., 2023). Within these circuits, the cortical amygdala (CoA; cortical amygdaloid nucleus) has recently emerged as a cortical node implicated in both binge-like and alcohol-dependent drinking (Ardinger et al., 2024; Kimbrough et al., 2020a; Roland et al., 2023; Xiao et al., 2024). However, this work has been conducted predominantly in male rodents, despite well-established sex differences in alcohol behavior and neural adaptation (Flores-Bonilla and Richardson, 2020; Kranzler et al., 2019), leaving it unclear whether CoA-dependent mechanisms generalize to females.

In addition, prior CoA studies have focused largely on alcohol consumption rather than withdrawal-associated affective states or large-scale circuit reorganization, leaving the contribution of the CoA to withdrawal-related behaviors and brain-wide network activity associated with withdrawal insufficiently defined. It is unclear whether alcohol dependence alters intrinsic CoA neuronal activity during withdrawal or whether CoA signaling contributes to the distributed connectivity disruptions characteristic of dependent states. Alcohol withdrawal is known to produce widespread alterations in neural activity across thalamic, cortical, and limbic circuits, suggesting that circuit-level reorganization may contribute to withdrawal-associated behavioral states (Becker, 2008; Kimbrough et al., 2020b; Koob, 2020; Koob and Volkow, 2016b). Since alcohol dependence engages distributed neural systems rather than isolated regions (Koob and Volkow, 2010, 2016b), understanding how the CoA influences large-scale connectivity is essential for identifying circuit mechanisms underlying alcohol drinking and withdrawal.

Here, we investigated the role of the CoA in regulating neuronal activity, alcohol-dependent drinking, withdrawal-related behaviors, and brain-wide connectivity during alcohol withdrawal in female mice. Specifically, we: (1) examined intrinsic CoA neuronal activity during withdrawal; (2) tested whether inhibition of the CoA in female mice significantly altered alcohol-dependent drinking behavior which previously observed in males (Roland et al., 2023); (3) assessed effects on withdrawal-related behaviors; and (4) determined whether CoA inhibition modifies brain-wide network connectivity during withdrawal. These studies address a critical sex-specific gap and define the contribution of the CoA to circuit-level mechanisms underlying alcohol dependence.

## Materials and Methods

### Animals

Group-housed eight-week-old female mice were obtained from Jackson Laboratories. Mice underwent a week of acclimation after delivery prior to experimental procedures. The mice were housed with a reverse 12-hour light/12-hour dark cycle on cage racks with 1/8’’ Bed-o-cobb bedding (Anderson, Maumee, OH, USA) and provided with over 8 g of nesting material for enrichment. Food (2018S Teklad from Envigo) and water were provided ad libitum. All experimental procedures were conducted in accordance with the guidelines of the National Institutes of Health (NIH) *Guide for the Care and Use of Laboratory Animals* and approved by the Purdue University Animal Care and Use Committee.

For electrophysiology, six female mice were used to evaluate the intrinsic neuronal excitability of CoA neurons. A total of 20 neurons were recorded from 2 groups, 3 mice/group (n = 10 cells; N = 3 animals/group). These mice were not included in the behavioral experiments and were used exclusively to assess neuronal excitability of the CoA.

Another separate cohort of mice was used for the behavioral testing and chemogenetic inhibition. After acclimation, mice were assigned to either an alcohol dependent group given CIE or AIR group. Groups were balanced based on their baseline drinking volumes, ensuring that each group had equivalent initial alcohol intake. Within each exposure group, mice were further subdivided to receive viral injections of either a control vector expressing mCherry or an inhibitory DREADD vector expressing hM4Di. This design resulted in four experimental groups: AIR::mCherry, AIR::hM4Di, CIE::mCherry, and CIE::hM4Di (AIR::mCherry, N = 9; AIR::hM4Di, N = 11; CIE::mCherry, N = 15; CIE::hM4Di, N = 24).

Following viral injection and recovery, mice underwent CIE or air exposure procedures as described below, followed by behavioral testing during withdrawal. For chemogenetic experiments during withdrawal, mice received intraperitoneal injections of saline or clozapine-N-oxide (CNO, 3 mg/kg; Tocris, Cat. No. 4936) before behavioral testing sessions.

### Electrophysiology

#### Brain slices preparation

Alcohol-naïve (AN) and Alcohol-dependence (AD) mice were anesthetized with ketamine (0.1 mg/g) and xylazine (0.16 mg/g) diluted in saline, and perfused within 1 min with ice-cold choline-based ACSF (in mM: 110 choline chloride, 30 NaHCO_3_, 25 dextrose, 11.6 ascorbic acid, 7 MgCl_2_, 3.1 Na-pyruvate, 2.5 KCl, 0.5 CaCl_2_; pH 7.3–7.4). Brains were rapidly removed and sectioned into 250 μm coronal slices using a vibratome (VT1000, Leica) in chilled choline ACSF. Slices were incubated at 32 °C in standard ACSF (in mM: 124 NaCl, 26 NaHCO_3_, 10 dextrose, 2.5 KCl, 2 CaCl_2_, 1.25 NaH_2_PO_4_, 0.8 MgCl_2_; pH 7.3–7.4) for 30 min and then equilibrated at room temperature for 2 h. All solutions were continuously bubbled with carbogen (95% O_2_/5% CO_2_), and slices remained viable for patch-clamp recordings for up to 10 h.

#### Whole-Cell Patch-Clamp Recordings

Recordings were performed in acute slices placed in a submerged chamber perfused with carbogenated ACSF (in mM: 124 NaCl, 26 NaHCO_3_, 10 dextrose, 2.5 KCl, 2 CaCl_2_, 1.25 NaH_2_PO_4_, 0.8 MgCl_2_; pH 7.3– 7.4). Neurons were visualized under infrared differential interference contrast optics (Olympus 4× and 40× water-immersion objectives, ROLERA™ Bolt camera). Patch electrodes (4–7 MΩ) were filled with internal solution containing (in mM): 130 K-gluconate, 5 KCl, 2 MgCl_2_, 10 HEPES, 2 MgATP, 0.3 Na_2_GTP, and 0.6 EGTA (pH 7.2–7.3; 292 mOsm). Signals were amplified with a Multiclamp 700B amplifier, digitized using a Digidata 1550, acquired in Clampex 10.4, and low-pass filtered at 20 kHz. Recordings were made in current-clamp mode with 0.6 s current steps (−40 to 150 pA, 10 pA increments). After baseline acquisition in ACSF, ethanol (190 Proof, Decon) was added to the bath to a final concentration of 88 mM, and responses were recorded for 10 min. No recordings were excluded from analysis. Data were analyzed using custom Python scripts.

#### Stereotaxic Surgery

For the chemogenetic study, mice were anesthetized using an isoflurane/oxygen mixture (1–3%) and secured in a stereotaxic apparatus (Injector: World Precision Instruments, UMP3T-2). Viral vectors containing either the inhibitory DREADD (AAV5-hSyn-hM4Di-mCherry; titer: 5 × 10^12^ vg/ml) or a reporter control (AAV5-hSyn-mCherry; titer: 5 × 10^12^ vg/ml) were obtained from Addgene (catalog no. 50475-AAV5 and 114472-AAV5). Eight-week-old female mice underwent stereotaxic injections of 350 nl of viral vectors bilaterally into the posterior cortical amygdala (CoA; coordinates: -2.9 AP, ±2.8 ML, -5.55 DV from bregma) using a microinjection syringe pump (World Precision Instruments, UMP3T-2) and 5 µl stereotaxic injection syringes (Hamilton, CAL 87943). The infusion rate was set at 70 nl/min for 5 min. Following infusion, the injectors were left in place for 8 min before removal. Following stereotaxic surgery, mice recovered in their home cages for 5–7 days with ad libitum access to food and water and no alcohol. After recovery, baseline alcohol drinking was assessed. The precise localization of injection sites and confirmation of viral infectivity were assessed post hoc using iDISCO immunolabeling imaging focused on the brain regions.

#### Chronic Intermittent Ethanol Vapor Exposure (Whole-Cell Patch-Clamp Recordings)

Mice were subjected to chronic intermittent ethanol (CIE) exposure in inhalation chambers (La Jolla Alcohol Research Inc., La Jolla, CA), as previously described (Becker and Hale, 1993; Becker and Lopez, 2004; Contet et al., 2011), During each CIE week, mice underwent four 16-hour periods of ethanol vapor intoxication, each separated by an 8-hour withdrawal period during which animals were exposed to air. For CIE weeks, the mice underwent four days of 16-hour intoxication followed by 8-hour abstinence (Monday to Friday), then 72-hour abstinence (Friday to Monday). Before exposure in the vapor chambers for CIE, mice were injected (i.p.) with an ethanol-pyrazole solution (1500 mg/kg EtOH and 68.1 mg/kg pyrazole) to induce initial ethanol intoxication, inhibit ethanol dehydrogenase, and block ethanol metabolism (Blomstrand et al., 1979; Feierman and Cederbaum, 1987). Blood ethanol concentrations were measured at the end of every CIE exposure period to ensure consistent ethanol intoxication across sessions. Air-exposed mice (AIR group) were handled identically and exposed to air. AIR mice received intraperitoneal injections of pyrazole in saline (68.1 mg/kg). Mice underwent six weeks of CIE or AIR exposure.

This continuous CIE paradigm was used exclusively for electrophysiological recordings to promote stable cellular and intrinsic membrane adaptations suitable for slice physiology.

### Behavioral Testing

Behavioral tests were conducted using a consistent testing framework across all experimental groups within a single defined withdrawal period. Behavioral assays were performed in the following fixed order across Withdrawal Days (WD): von Frey testing on WD3–WD4, Open Field Test on WD5, and Tail Suspension Test on WD6. Tissue collection occurred on WD7 (Figure 1A). Groups of mice received saline and CNO injections (3 mg/kg) on counterbalanced testing days to control for injection- and order-related effects.

**Figure 1.**
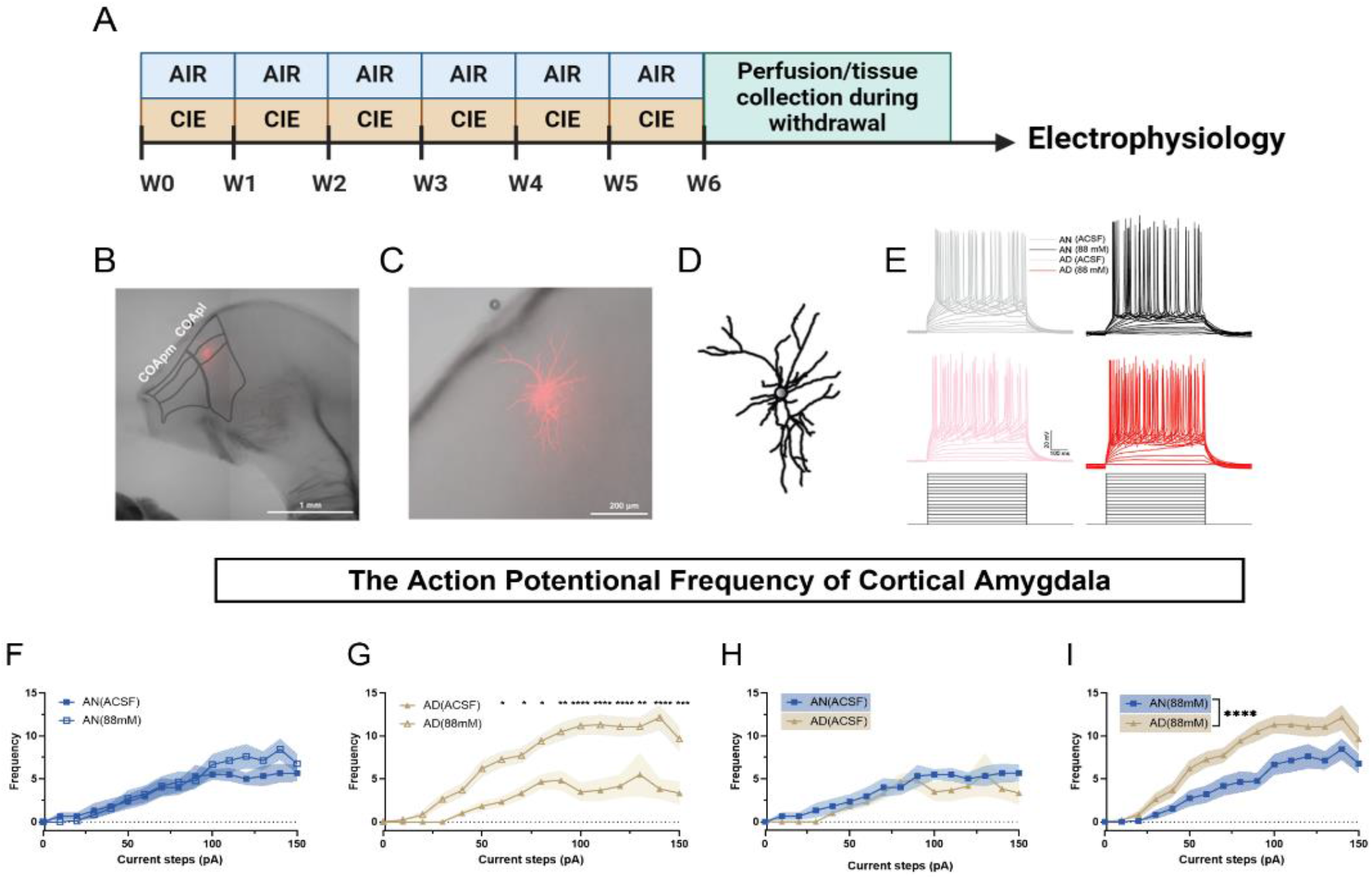
Intrinsic activity of cortical amygdala neurons during alcohol withdrawal. **(A)** Experimental timeline for electrophysiological recordings. **(B-C)** Confocal images of biocytin-filled patched neurons. Scale bars: 1 mm, 200 μm. Reconstructed neuron morphology. Scale bar: 100 μm. Representative firing traces from alcohol-naïve (AN) and alcohol-dependent (AD) CoA neurons under ACSF or acute alcohol (88 mM). **(F–I)** Firing frequency during step current injections (0–150 pA). Acute alcohol did not affect AN neurons (F; interaction F(15,352) = 0.6415, P = 0.8401) but increased firing in AD neurons (G; interaction F(15,352) = 3.232, P < 0.0001). Baseline firing was unchanged (H), while firing under alcohol was higher in AD neurons (I; main effect: F(1,416) = 77.08, P < 0.0001). Data: mean ± SEM; AN (n = 14 neurons, N = 3 mice); AD (n = 14 neurons, N = 3 mice).

#### Two-bottle choice / Chronic intermittent ethanol vapor exposure

To model voluntary alcohol consumption and dependence-associated behaviors, we employed the well-established two-bottle choice combined with chronic intermittent ethanol vapor exposure (2BC/CIE) paradigm (Becker and Lopez, 2004; Borgonetti et al., 2023; Varodayan et al., 2023; Xiao et al., 2023). In this paradigm, weeks of voluntary alcohol consumption via 2BC sessions alternated with weeks of CIE exposure, forming a repeating cycle. All vapor parameters, pyrazole/saline administration, blood alcohol concentration monitoring, and chamber conditions were identical to those described above.

During 2BC weeks, mice were provided with two bottles, one containing water and the other alcohol (15% vol/vol; Decon), for five consecutive days for 2 hours starting at the beginning of the dark phase of their circadian cycle. Prior to the beginning of the 2BC/CIE paradigm, the mice underwent 2 weeks of baseline 2BC testing (BSL; weeks 1 to 2) and were then divided into AIR and CIE groups with equivalent baseline intake. Subsequently, the mice underwent six cycles of 2BC/CIE (weeks 3 to 13), followed by an additional week of CIE exposure (week 14) prior to testing of withdrawal behavior and brain tissue collection (week 15).

#### Chemogenetic Manipulation During Final 2BC Week

During the final 2BC testing week (week 13), mice were tested to evaluate the effect of chemogenetic inhibition on drinking behavior. On Testing Day 1, mice underwent a 2BC session without injection. On Testing Days 2 and 4, mice received saline injections (vehicle, i.p.) 30 minutes before 2BC. On Testing Days 3 and 5, mice received CNO injections (3 mg/kg, i.p.) 30 minutes before 2BC. Alcohol intake on CNO days was compared with saline and non-injection days to determine the effect of CoA inhibition on voluntary drinking.

#### von Frey

Mechanical pain sensitivity was assessed using the von Frey assay, a well-established method for evaluating nociceptive alterations during alcohol withdrawal (Brandner et al., 2023; Deuis et al., 2017). Mechanical pain sensitivity was assessed at baseline (Week 2) and again during alcohol withdrawal on Withdrawal Days 3–4 (WD3–WD4, week 15). During withdrawal testing, mice received an intraperitoneal injection of either saline or CNO 30 minutes prior to testing, with treatment order counterbalanced within subjects across days.

For the von Frey test, mice were individually placed in a grid with a metal mesh floor and habituated for 30 minutes before testing. Mechanical sensitivity was assessed by an electronic von Frey apparatus (BioSeb Lab Instruments, Vitrolles Cedex, France). A rigid filament connected to a handheld force transducer was applied vertically to the plantar surface of each hind paw (left and right) from below, with force gradually increased until paw withdrawal occurred. The maximum force required to elicit withdrawal (grams) and interaction time (seconds) were automatically recorded. Each hind paw was tested three times, and the mean of six measurements (left and right paws) was used as the mechanical withdrawal threshold for each animal.

#### Open Field Test

The Open Field Test was utilized to assess anxiety-like behavior during abstinence from alcohol following final CIE exposure (Quijano Cardé and De Biasi, 2022). On Withdrawal Day 5 (WD5; week 15), mice were placed in a 40 × 40 × 40 cm box (Maze Engineers, Boston, MA) for 10 minutes. The center zone was defined as a 20 × 20 cm square. Locomotor and anxiety-related measures, including total distance traveled, time spent in the center, and number of center entries, were automatically recorded using EthoVision XT tracking software (Noldus, Leesburg, VA). Mice received an intraperitoneal injection of either saline or CNO 30 minutes prior to testing. The treatment order was counterbalanced across experimental groups to control for injection-related effects.

#### Tail Suspension Test

Depressive-like behavior was assessed using the tail suspension test (Cryan et al., 2005; Koob and Le Moal, 2008). On withdrawal day 6 (WD6; week 15), mice were suspended by the tail for 6 minutes in a tail suspension apparatus (BIO-TST5, Bioseb, USA), and immobility time (s) was automatically quantified using integrated strain sensors. Mice received an intraperitoneal injection of either saline or CNO 30 minutes prior to testing. Treatment assignment was counterbalanced across groups.

#### Tissue Collection

On withdrawal day 7 (WD7, Week 15), mice received a final intraperitoneal injection of saline/ CNO at a time corresponding to a typical 2BC session. Two hours after injection, a time point chosen to coincide with peak c-Fos expression following CNO administration, mice were deeply anesthetized and perfused with 20 mL of 0.9% heparinized saline followed by 20 mL of 4% paraformaldehyde (PFA) in phosphate-buffered saline (PBS). Brains were immediately extracted, post-fixed in 4% PFA at 4 °C for 24 hours, and then transferred to 1× PBS for long term storage prior to iDISCO+ processing.

#### iDISCO+

We examined whole-brain immunostaining for protein from the immediate early gene *c-Fos* as a marker representative of neuronal activity following our established protocols (Ardinger et al., 2024; Kimbrough et al., 2021; Kimbrough et al., 2020a), as outlined in the iDISCO protocol (https://idisco.info/idisco-protocol/). Briefly, whole brains were dehydrated in a series of methanol concentrations (20%, 40%, 60%, 80%, and 100%) in double-distilled water (ddH2O) for 1 hour at each concentration. Samples were then bleached overnight in 5% hydrogen peroxide in methanol and subsequently rehydrated in a reverse methanol series (100%-20%) in ddH2O. Next, the brains were washed in PBS with 0.2% Triton X-100, and permeabilized for 2 days at 37°C in a solution containing 0.3 M glycine and 20% DMSO in PBS with 0.2% Triton X-100. Tissue was then blocked for 2 days at 37°C in PBS with 0.2% Triton X-100, 10% DMSO, and 6% donkey serum (Jackson ImmunoResearch, #AB_2337258). Primary antibodies (rabbit anti-cFos, 1:1000, Synaptic Systems #226-003; chicken anti-mCherry, 1:1000, Aves Labs MCHERRY-0020) were incubated for 7 days in PBS with 0.2% Tween-20, 10% DMSO, 6% donkey serum, and 10 µg/mL heparin. Following this, the brains were washed in PBS with 0.2% Tween-20 and heparin, then incubated for 7 days in a secondary antibody solution containing donkey anti-rabbit Alexa Fluor 647 (1:500; Thermo Fisher Scientific #A-31573) and donkey anti-chicken Alexa Fluor 568 (1:500; Thermo Fisher Scientific #A-78950) in PBS with 0.2% Tween-20, 10% DMSO, 5% donkey serum, and heparin. After additional washes in PBS with 0.2% Tween-20 and heparin, the brains were incubated in an increasing series of methanol concentrations (20%-100%) in PBS. The following day, the brains were incubated in a solution of 66% dichloromethane and 33% methanol for 3 hours, followed by two consecutive 15-minute incubations in 100% dichloromethane. Finally, brains were transferred to dibenzyl ether for refractive index matching and stored until imaging.

#### Image Acquisition

Cleared brain hemispheres were imaged in the sagittal orientation using a light sheet microscope (Ultramicroscope II, Miltenyi Biotec, Bielefeld, Germany) equipped with an Olympus MVPLAPO 2X/0.5 objective. The right hemisphere of the brain was imaged, and image acquisition was controlled using ImspectorPro software. Imaging parameters included a light-sheet numerical aperture of 0.026, a sheet width of 100 %, exposure time of 150 ms, and 3 µm Z-step resolution.

mCherry fluorescence was acquired in the 568 nm channel to visualize viral expression, autofluorescence was collected in the 488 nm channel for anatomical registration, and Fos immunolabeling was imaged in the 640 nm channel. Image volumes were imported into the ClearMap analysis pipeline for atlas registration, cell detection, and regional quantification.

#### ClearMap Analysis

Images were processed using ClearMap v2.0 on an Ubuntu operating system following the validated ClearMap 2.1 pipeline(Kirst et al., 2020; Renier et al., 2016; Renier et al., 2014), Automated cell detection parameters were optimized based on signal intensity and cellular morphology. Three-dimensional affine registration aligned the 488 nm autofluorescence channel to the Allen Brain Atlas, after which the 640 nm c-Fos channel was registered to the aligned reference space. Fos-positive (Fos^+^) nuclei were detected using a ClearMap, and each dataset was visually inspected to verify registration accuracy and prevent double-counting artifacts.

Region-specific Fos^+^ cell counts were extracted at 3 µm spatial resolution across annotated brain structures. These counts served as the quantitative basis for subsequent normalization, correlation, and network analyses. Representative images of cleared brain hemispheres and Fos-labeled nuclei are shown in Figure S2. Summary counts from selected brain regions, along with their broader anatomical classifications (e.g., isocortex, thalamus, hypothalamus, midbrain, and hindbrain, as defined by the Allen Brain Atlas), are presented in Table S1. Region-specific Fos^+^ counts were subsequently used for both correlation and graph-theory analyses described below.

#### Correlations Analysis between Brain Regions

For our analysis, to identify the impact of inhibiting the CoA during alcohol dependence, we focused on brains from mice that were alcohol dependent (CIE), with active virus (hM4Di), that were given either saline or CNO prior to perfusion. From these brains, Fos^+^ cell counts were log^10^-transformed to reduce right-skew and heteroscedasticity, consistent with prior whole-brain immediate early gene mapping and ClearMap-based analyses (Ardinger et al., 2024; Kimbrough et al., 2021; Kimbrough et al., 2020b; Roland et al., 2023). Pearson correlations between each brain region examined were calculated for each treatment (CIE::hM4Di::Saline and CIE::hM4Di::CNO). Value transformed correlations and heatmaps were generated in R.

#### Graph Theory Identification of Betweenness and Degree Networks

A graph theory-based approach was employed to identify the functional neural networks associated with the CNO/saline treatment of alcohol dependent mice. Using this approach, we constructed network maps and identified key nodes (i.e., brain regions) involved in connectivity patterns. For each network, we computed degree and betweenness centrality values to quantify the relative importance of individual brain regions. Definitions and applications of degree and betweenness centrality follow established graph-theoretical frameworks for brain networks (Bullmore and Sporns, 2009), and have been widely applied to mouse whole-brain Fos connectivity analyses (Wheeler et al., 2013). Degree represents the number of direct functional connections to a brain region, while betweenness quantifies how often a brain region lies on the shortest path of functional connections between other brain regions (Freeman, 1978; Rubinov and Sporns, 2010a).

Degree centrality was defined as:

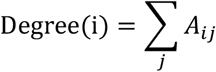

where *A*_*ij*_ represents the adjacency or weighted connectivity matrix between region *i* and region *j*. Betweenness centrality calculated as (Rubinov and Sporns, 2010a; Sporns et al., 2007):

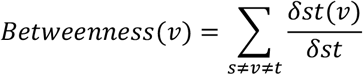

where *σst* is the total number of shortest paths between nodes s and t, and *σst*(*v*) represents the subset of those paths that pass-through node *v*.

Regions ranking in the top 15 brain regions (top 10% of all brain areas) for either metric were classified as hub regions, as they are likely to play central roles in maintaining network structure and facilitating communication across the brain. These metrics were computed using a customized Python library (https://github.com/aestrivex/bctpy), adapted from the MATLAB-based Brain Connectivity Toolbox (Rubinov and Sporns, 2010b). Network visualization was carried out in Gephi 0.10.1(Bastian et al., 2009) with the Force Atlas 2 algorithm (Jacomy et al., 2014), with selected brain regions manually repositioned for visual clarity. Final figures were prepared in Adobe Illustrator for presentation.

#### Statistical Analysis

Data analysis and curve fitting were performed using R, GraphPad Prism 9, Imaris, and ImageJ. Behavioral and electrophysiological data were analyzed using GraphPad Prism, whereas whole-brain imaging and connectivity analyses were performed using Imaris, Python and R.

Single comparisons between two groups were conducted using either a paired or unpaired two-tailed Student’s t-test, as appropriate. For comparisons involving multiple factors, two-way ANOVA, three-way ANOVA, or two-way repeated-measures (RM) ANOVA were used depending on the experimental design. Two-way RM ANOVA was applied when repeated measurements were obtained from the same subjects. Mixed-effects models were used when repeated-measures datasets contained missing values or unequal sample sizes, allowing inclusion of all available observations without requiring balanced data. When significant main effects or interactions were detected, Sidak’s multiple-comparisons tests were used for post hoc analyses. The number of experimental samples (N) in each group is specified in the corresponding figure legends. Data are presented as mean ± standard error of the mean (SEM). Statistical significance was defined as P < 0.05 (*), P < 0.01 (**), and P < 0.001 (***).

## Results

### 1. Chronic Intermittent Ethanol Vapor Exposure Alters Ethanol Sensitivity of Cortical Amygdala Neurons

To investigate the impact of alcohol dependence on neuronal activity in the cortical amygdala (CoA), we employed the continued chronic intermittent ethanol vapor exposure (CIE) model in mice, a well-established approach for inducing alcohol dependence followed by a one-week withdrawal period before brain tissue was collected for electrophysiological analysis of intrinsic neuronal properties (Becker and Lopez, 2004; Kirson et al., 2021; Nimitvilai et al., 2016) (Figure 1A-E). Whole-cell current-clamp recordings were obtained from CoA neurons in acute brain slices to evaluate intrinsic membrane properties and firing responses under baseline artificial cerebrospinal fluid (ACSF) conditions and during acute ethanol application (88 mM).

Under baseline conditions with artificial cerebrospinal fluid (ACSF), neuronal firing rates within the CoA did not significantly differ between the alcohol-dependent (AD) and AN groups (Figure 1H), indicating that chronic CIE exposure did not alter baseline firing output during withdrawal. Consistent with this observation, acute alcohol bath application (88 mM) to brain slices from alcohol-naive (AN) mice did not significantly alter neuronal firing across current steps (Figure 1F), suggesting minimal acute alcohol sensitivity in naive neurons.

In contrast, CoA neurons from AD mice exhibited enhanced responsiveness to acute alcohol. Application of 88 mM alcohol significantly increased firing frequency across current steps compared with ACSF within the AD group (Figure 1G). Two-way repeated-measures ANOVA revealed a significant CIE × current step interaction (F(15,352) = 3.232, P < 0.0001). Post hoc Sidak comparisons demonstrated elevated firing in AD neurons under 88 mM alcohol relative to ACSF at multiple current steps (60–150 pA). Consistently, firing frequencies under acute alcohol were significantly greater in AD compared with AN slices (Figure 1I; main effect of CIE: F(1,416) = 77.08, P < 0.0001), indicating that chronic alcohol exposure enhances neuronal sensitivity to subsequent alcohol challenge.

To further assess intrinsic membrane properties, input resistance was measured under ACSF conditions. Neurons from AD mice displayed significantly higher input resistance compared to those from AN mice across current steps (Figure 1L; two-way repeated-measures ANOVA, CIE treatment × current step interaction: F(19,360) = 4.196, P < 0.0001). Notably, this increase in input resistance did not translate into higher baseline firing frequency, indicating a dissociation between intrinsic resistance and action potential output under ACSF conditions.

Notably, acute alcohol exposure differentially influenced intrinsic properties between groups. While acute alcohol had no effect on input resistance in AN neurons (Figure 1J), neurons from AD mice exhibited a significant alcohol-induced change in input resistance (Figure 1K; two-way repeated-measures ANOVA, main effect of alcohol: F(1,520) = 6.229, P = 0.0129). Under 88 mM alcohol conditions, group differences in firing frequency across current steps were attenuated (Figure 1M; two-way repeated-measures ANOVA, CIE treatment × current step interaction: F(19,520) = 2.672, P = 0.0002). Post hoc Sidak’s multiple comparisons showed that, within the AD group, firing frequency under 88 mM alcohol was significantly higher than under ACSF at only one current step (90 pA), indicating a reduced excitability contrast across conditions. This interpretation is further supported by input resistance measurements: AN neurons showed no change in input resistance between ACSF and alcohol conditions (Figure 1J), whereas AD neurons exhibited a significant alcohol-induced change (Figure 1K; main effect of alcohol: F(1,520) = 6.229, P = 0.0129). Together, these findings suggest that acute alcohol alters intrinsic membrane properties in CoA neurons from alcohol-dependent mice, reshaping excitability–output coupling rather than restoring baseline firing levels.

Collectively, chronic alcohol exposure alters intrinsic membrane properties of CoA neurons without elevating baseline firing, whereas acute alcohol selectively increases firing in alcohol-dependent neurons. These results demonstrate enhanced alcohol sensitivity rather than normalization of excitability and suggest a cellular substrate for escalated drinking and relapse risk.

### 2. Chemogenetic inhibition of the Cortical Amygdala reduces alcohol consumption in alcohol-dependent mice

Given that CoA neurons in alcohol-dependent (AD) mice exhibited increased sensitivity to alcohol, we hypothesized that the CoA plays a role in alcohol dependence-related behaviors. To test this hypothesis, we used designer receptors exclusively activated by designer drugs (DREADDs) to selectively inhibit CoA activity during the final weeks of the 2BC/CIE paradigm (Figure 2A-B).

**Figure 2.**
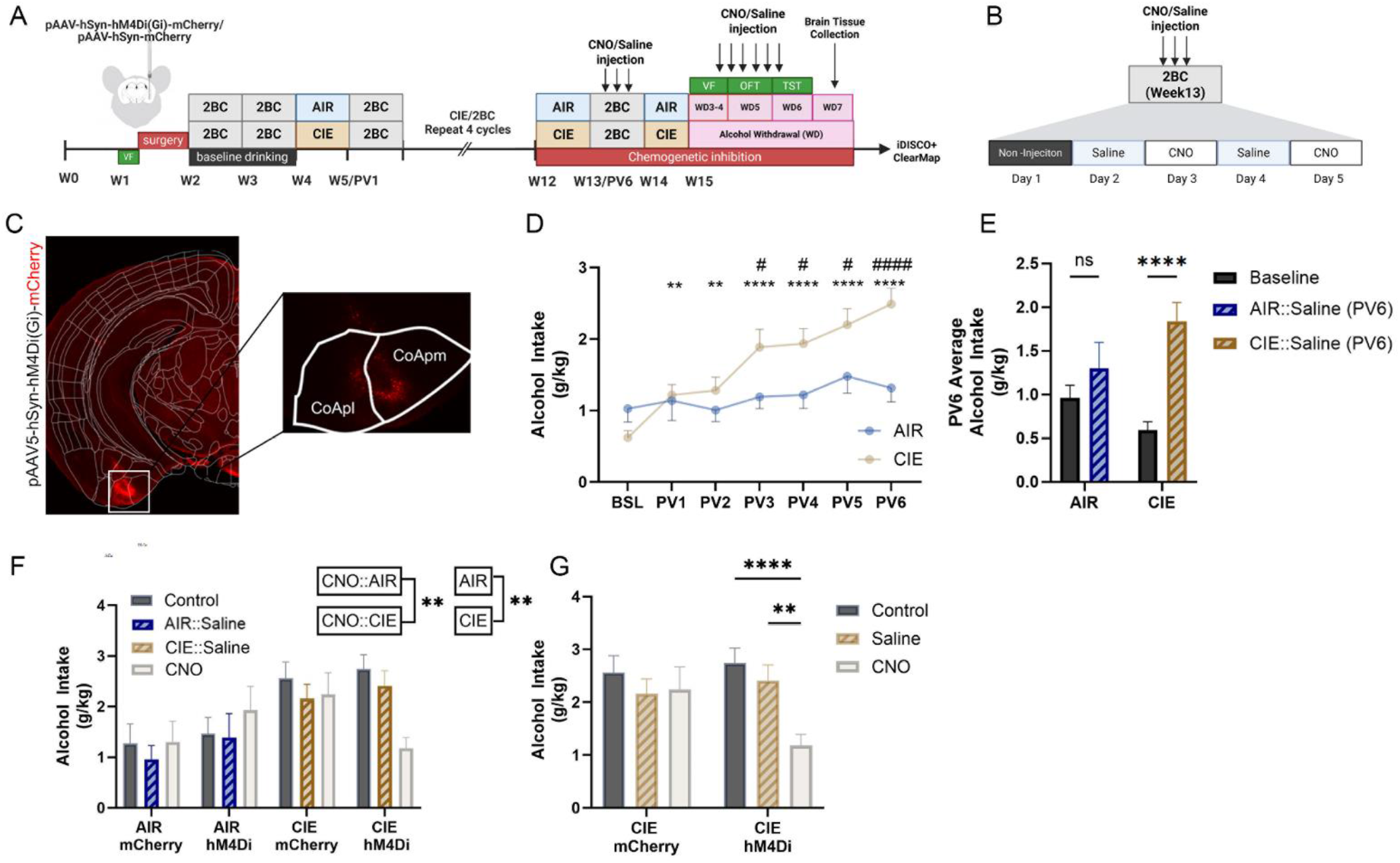
Chronic intermittent ethanol exposure escalates alcohol intake and chemogenetic inhibition of the cortical amygdala reduces drinking in dependent mice. **(A)** Experimental timeline for two-bottle choice (2BC) drinking paired with chronic intermittent ethanol vapor exposure (CIE). Withdrawal tests: von Frey (WD3–4), open field (WD5), tail suspension (WD6); brains collected WD7. **(B)** Experimental design for chemogenetic manipulation during the final 2BC week. **(C)** Viral targeting of the CoA. **(D)** Weekly alcohol intake across 2BC/CIE cycles (two-way mixed-effects ANOVA, interaction F(6,455) = 4.629, P = 0.0001; Sidak’s multiple-comparisons test *P < 0.05, **P < 0.01, ****P < 0.0001). **(E)** Alcohol intake during the final week (two-way mixed-effects ANOVA, interaction F(1,69) = 6.949, P = 0.0103; Sidak’s multiple-comparisons test ***P < 0.0001). **(F)** CNO reduced alcohol intake in CIE::hM4Di mice but not controls (three-way ANOVA, interaction F(2,110) = 5.575, P = 0.0049; AIR vs. CIE: F(1, 55) = 9.483, P = 0.0032). **(G)** Within-group comparisons confirmed reduced intake in CIE::hM4Di mice (mixed-effects model with Sidak test **P = 0.01, ****P < 0.0001). Data: mean ± SEM; CIE N = 44, AIR N = 28; AIR::mCherry N = 9; AIR::hM4Di N = 11; CIE::mCherry N = 15; CIE::hM4Di N = 23.

To evaluate the escalation of alcohol intake over time, we performed a two-way repeated-measures ANOVA with time (baseline drinking and post-vapor weeks) and alcohol treatment (AIR and CIE) as factors (Figure 2D). This analysis revealed a significant time × treatment interaction (F(6, 455) = 4.629, P < 0.0001). Post hoc Sidak’s multiple-comparisons tests indicated that CIE mice consumed significantly more alcohol than both AIR controls and their own baseline during post-vapor weeks 3 through 6. To further validate this escalation, we conducted a focused two-way repeated-measures ANOVA comparing only baseline and post-vapor week 6 (PV6) (Figure 2E), which also revealed a significant time × treatment interaction (F(1, 148) = 5.525, P = 0.0201). Post hoc Sidak’s multiple-comparisons tests confirmed that alcohol intake at PV6 was significantly elevated in CIE mice relative to both AIR mice and their own baseline, supporting the conclusion that CIE exposure leads to persistent increases in alcohol intake.

During the final week of 2BC drinking, CNO administration reduced alcohol intake in CIE mice with CoA inhibition. A three-way ANOVA revealed a significant interaction between CNO treatment and alcohol exposure (AIR vs. CIE) (Figure 2F; CNO × CIE interaction: F(6, 112) = 4.142, P = 0.0009), along with a main effect of CIE treatment (F(3, 56) = 2.997, P = 0.0382), indicating that the effect of CNO depended on dependence status. To further examine this interaction, we conducted a two-way repeated-measures ANOVA within the CIE group, which revealed a significant interaction between virus and CNO treatment (Figure 2G; F(2, 76) = 3.78, P = 0.0272). Post hoc Sidak’s multiple-comparisons tests showed that CNO significantly reduced alcohol intake in CIE mice expressing hM4Di in the CoA, compared with both saline-injected (P = 0.0010) and non-injected (P < 0.0001) conditions. No significant differences were observed in CIE mice expressing the control mCherry virus.

These findings demonstrate that chemogenetic inhibition of the CoA reduces alcohol intake in dependent female mice, supporting a causal role for CoA activity in driving excessive alcohol consumption. Electrophysiological validation confirmed that CNO effectively suppressed activity in hM4Di-expressing CoA neurons (Figure S1A–C).

### 3. Chemogenetic inhibition of the cortical amygdala does not alter withdrawal-associated behavioral phenotypes

To determine whether CoA inhibition influences behavioral signs of alcohol withdrawal, we assessed mechanical sensitivity, anxiety-like behavior, and depressive-like behavior during withdrawal. Hyperalgesia was measured using the von Frey test at baseline and two withdrawal time points following saline or CNO (counterbalanced; Figure 3A–D). CoA inhibition did not alter withdrawal-induced hyperalgesia, as no virus × drug interaction was detected. However, a two-way repeated-measures ANOVA showed a significant effect of alcohol exposure (CIE vs AIR; F(1,38)=13.38, P=0.0008). Post hoc analysis confirmed reduced mechanical thresholds in CIE mice compared with AIR controls and baseline (P<0.0001), indicating withdrawal-associated hyperalgesia. These results confirm that CIE exposure leads to withdrawal-associated hyperalgesia that is not significantly modulated by chemogenetic inhibition of the CoA.

**Figure 3.**
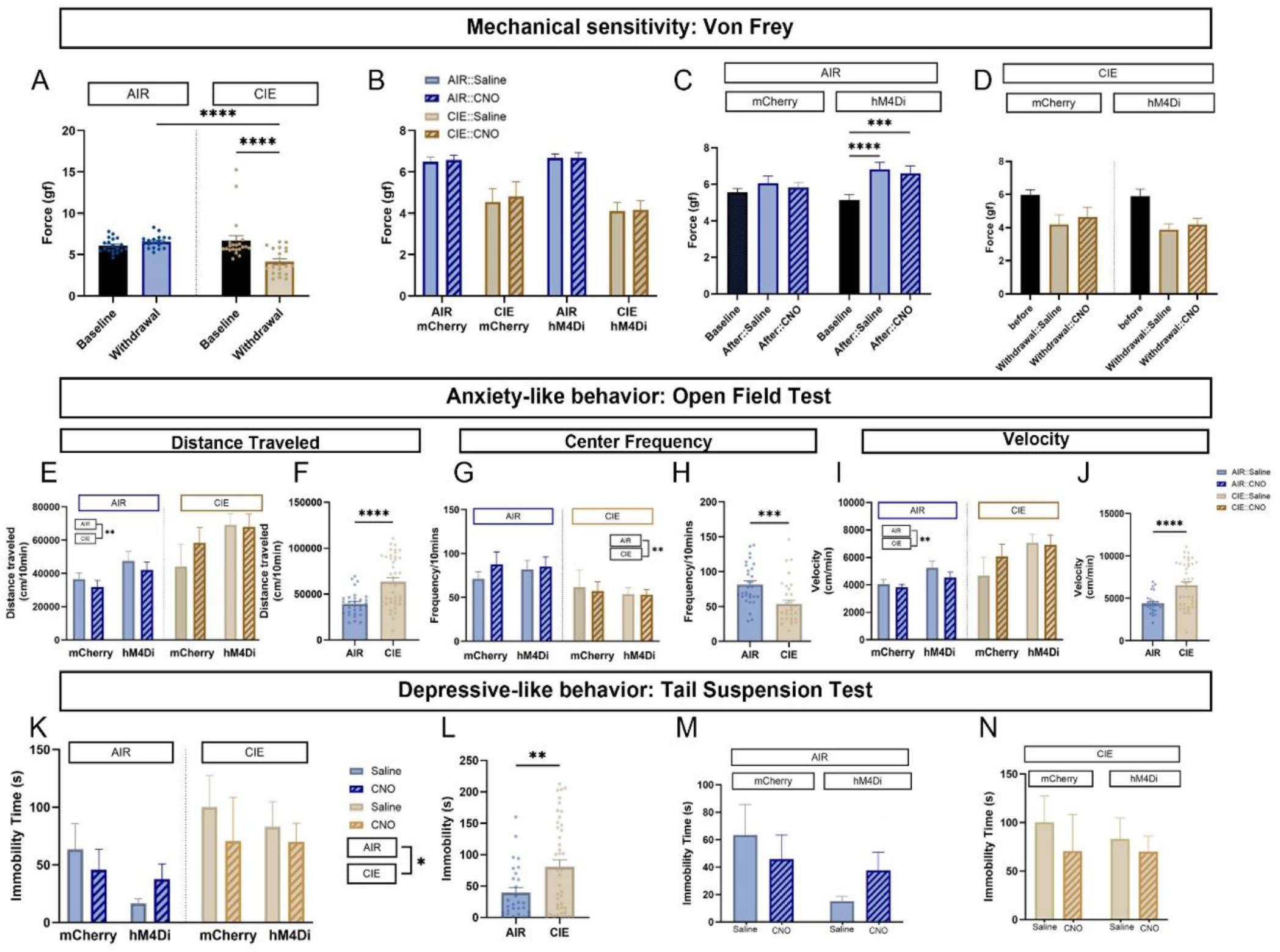
Chemogenetic inhibition of the cortical amygdala does not alter withdrawal-associated behavior. **(A-D)** Mechanical sensitivity assessed by von Frey. (A) Withdrawal thresholds at baseline and WD3–WD4 in AIR (blue) and CIE (gold) mice (interaction F(1,38) = 13.38, P = 0.0008; ***P < 0.0001). (B) Group comparisons across virus and drug during withdrawal (CIE × virus interaction: F(1,72) = 45.17, P < 0.0001). **(C–D)** Within-group analyses in AIR and CIE mice (RM two-way ANOVA with Sidak’s multiple-comparisons test). **(E–J)** Open field test during withdrawal. (E, G, I) Distance traveled, center frequency, and velocity across groups; no effects of CoA inhibition (three-way ANOVA). (F, H, J) Collapsed AIR vs CIE comparisons show increased distance and velocity and reduced center frequency in CIE mice (**P < 0.001, ***P < 0.0001). **(K–N)** Tail suspension test. (K) Immobility across groups (three-way mixed-effects ANOVA, interaction n.s.; main effect of alcohol exposure F(1,64) = 3.996, P = 0.0499). (L) Increased immobility in CIE vs AIR (**P < 0.01). (M–N) Within-group analyses (two-way ANOVA; interaction n.s.). Data are mean ± SEM. Group sizes as in Figure 2.

Following the assessment of pain sensitivity, we examined whether inhibition of the CoA influenced alcohol withdrawal-related increases in anxiety-like behavior using the open field test. Mice expressing either hM4Di or a mCherry virus received either CNO or saline injections 30 minutes prior to testing three days into withdrawal. Across all treatment conditions (CIE × Virus × Drug), a three-way repeated-measurement ANOVA revealed no significant effects of CNO injection or viral vector on open field behavior (Figure 3E, G, I). However, consistent differences were observed between AIR and CIE mice. CIE mice showed increased distance traveled (Figure 3F, F(1,14)=19.59, P=0.006) and velocity (Figure 3J, F(1,64)=11.67, P=0.0011), along with reduced center frequency (Figure 3H, F(1,64)=11.37, P=0.0013), consistent with increased anxiety-like behavior. These results demonstrate that the mice showed increased anxiety-like behavior during withdrawal from alcohol dependence, but that inhibition of the CoA did not impact this behavior. Furthermore, no significant differences in distance traveled were observed between CIE::mCherry and CIE::hM4Di groups following either saline or CNO administration (Figure 3E), indicating that the reduction in alcohol intake was not due to CNO treatment or locomotor impairment.

We subsequently investigated the effect of inhibiting the CoA on alcohol withdrawal-induced depression-like behaviors using the tail suspension test, where longer immobility times are indicative of greater depression-like responses (Figure 3K–N). Three-way ANOVA revealed no significant effects of viral manipulation or CNO treatment on immobility time. However, a significant main effect of alcohol exposure history was detected (Figure 3L, AIR VS. CIE: F (1, 64) = 3.996, P=0.0499). To further evaluate treatment effects within each exposure group, two-way ANOVAs were conducted separately for AIR and CIE mice. No significant effects of viral expression or CNO treatment were observed in either group (Figures 3M-N).

Given there was no virus or drug specific effect between groups, we combined the mCherry and hM4Di groups and conducted an unpaired t-test comparing AIR and CIE groups. Alcohol dependent CIE mice showed a significant increase in immobility time compared with alcohol non-dependent AIR mice (Figure 3L, AIR vs.CIE: mean± SEM: 40.91 ± 15.26; t = 2.68, P=0.0092), showing heightened depression-like behavior during withdrawal.

### 4. Inhibition of the Cortical Amygdala during withdrawal from alcohol dependence results in widespread neuronal inactivation

To determine how CoA inhibition affects brain activity during alcohol withdrawal, alcohol-dependent (CIE) mice received CNO or saline during withdrawal, and brains were collected for Fos detaction across mulptiple regions. Unpaired t-tests showed that CNO-treated group had significantly reduced Fos expression in both the anterior and posterior CoA subdivisions (Figure 4C; CoAa: P = 0.03; CoAp: P = 0.0076), confirming effective chemogenetic suppression. To further analyze Fos counts, values were log_10_transformed before statistical testing. unpaired t-test of the transformed counts showed that 64 of 80 analyzed brain regions exhibited decreased Fos expression (Figure 4D; P < 0.05), indicating that CoA inhibition broadly reduced neuronal activity in alcohol dependent mice.

**Figure 4.**
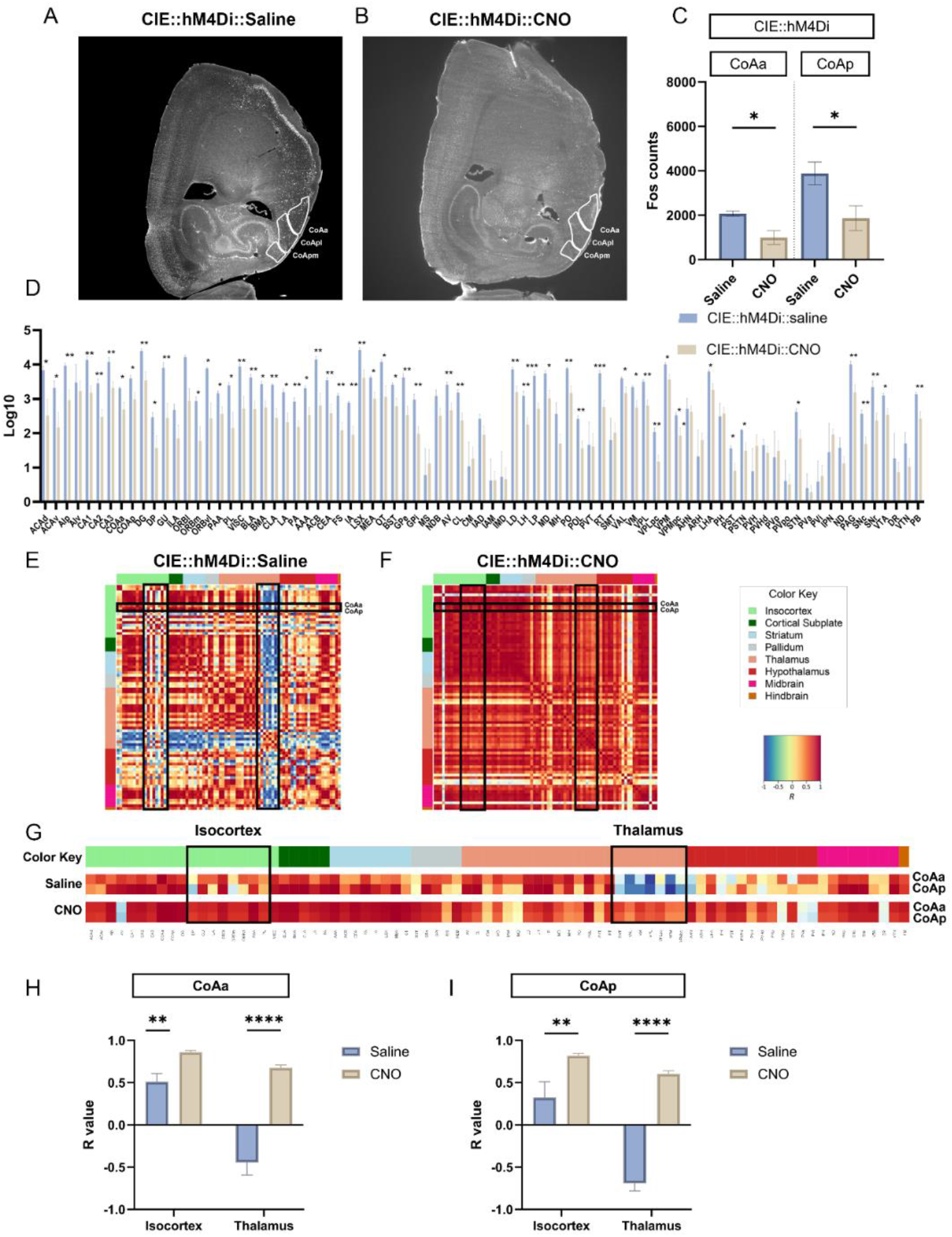
Chemogenetic inhibition of the cortical amygdala alters neuronal activity and functional connectivity during alcohol withdrawal. **(A-B)** Cleared brain hemispheres from CIE::hM4Di mice following saline or CNO. **(C)** Fos-positive cells in anterior (CoAa) and posterior (CoAp) CoA following saline or CNO (two-way RM ANOVA with Sidak test *P < 0.05). **(D)** Log_10_-transformed Fos counts across brain regions showing reduced activation after CNO (unpaired t-test, *P < 0.05, **P < 0.01, ***P < 0.001). **(E-F)** Whole-brain correlation matrices under saline and CNO. **(G)** Enlarged CoA correlation profiles highlighting the isocortex and thalamus connectivity changes. **(H-I)** Increased CoA connectivity with the isocortex and thalamus regions after inhibition (unpaired t-test **P < 0.01, ***P < 0.0001). Data are mean ± SEM. Full names of abbreviations are provided in Table S1, Log_10_-transformed Fos counts are provided in Table S3.

Pearson correlation analysis was then performed on the log_10_-transformed Fos values, and the resulting correlation matrices were visualized as heatmaps to evaluate brain-wide co-activation patterns (Figure 4E–G). To further define how CoA inhibition altered CoA-related network activity during alcohol withdrawal, we focused on two major clusters identified in the heatmaps: the isocortex and the thalamus (Figure 4H–I). Overall, CoA inhibition shifted CoAa–thalamus and CoAp–thalamus correlations from negative to positive values, indicating a marked increase in coordinated activity within thalamic-associated networks. Unpaired t-tests of the correlation coefficients (R values) showed significantly greater shifts in CoAa-related correlations with both the isocortex (Figure 4H; t(7) = 3.933, P = 0.0039) and thalamus (t(6) = 8.098, P = 0.0002) after CoA inhibition. Similarly, CoAp-related correlations with the isocortex (Figure 4I; t(7) = 2.938, P = 0.0201) and thalamus (t(6) = 15.83, P < 0.0001) were also significantly altered following CoA inhibition. These findings suggest that inhibiting the CoA suppresses local CoA activity while enhancing coordinated CoA–isocortex and CoA–thalamus co-activation during alcohol withdrawal relative to the non-inhibited condition.

### 5. Identification of key brain regions associated with Cortical Amygdala function during withdrawal from alcohol

After completing the whole-brain modular analysis, we next applied network analysis to examine node-level brain region properties in CIE::hM4Di::Saline and CIE::hM4Di::CNO mice (Figure 5A–B). In this analysis, degree centrality, shown by node color, reflects the number of direct connections a brain region has with other regions, whereas betweenness centrality, shown by node size, reflects how often a region lies on the shortest paths between other nodes and therefore serves as a bridge within the network.

**Figure 5.**
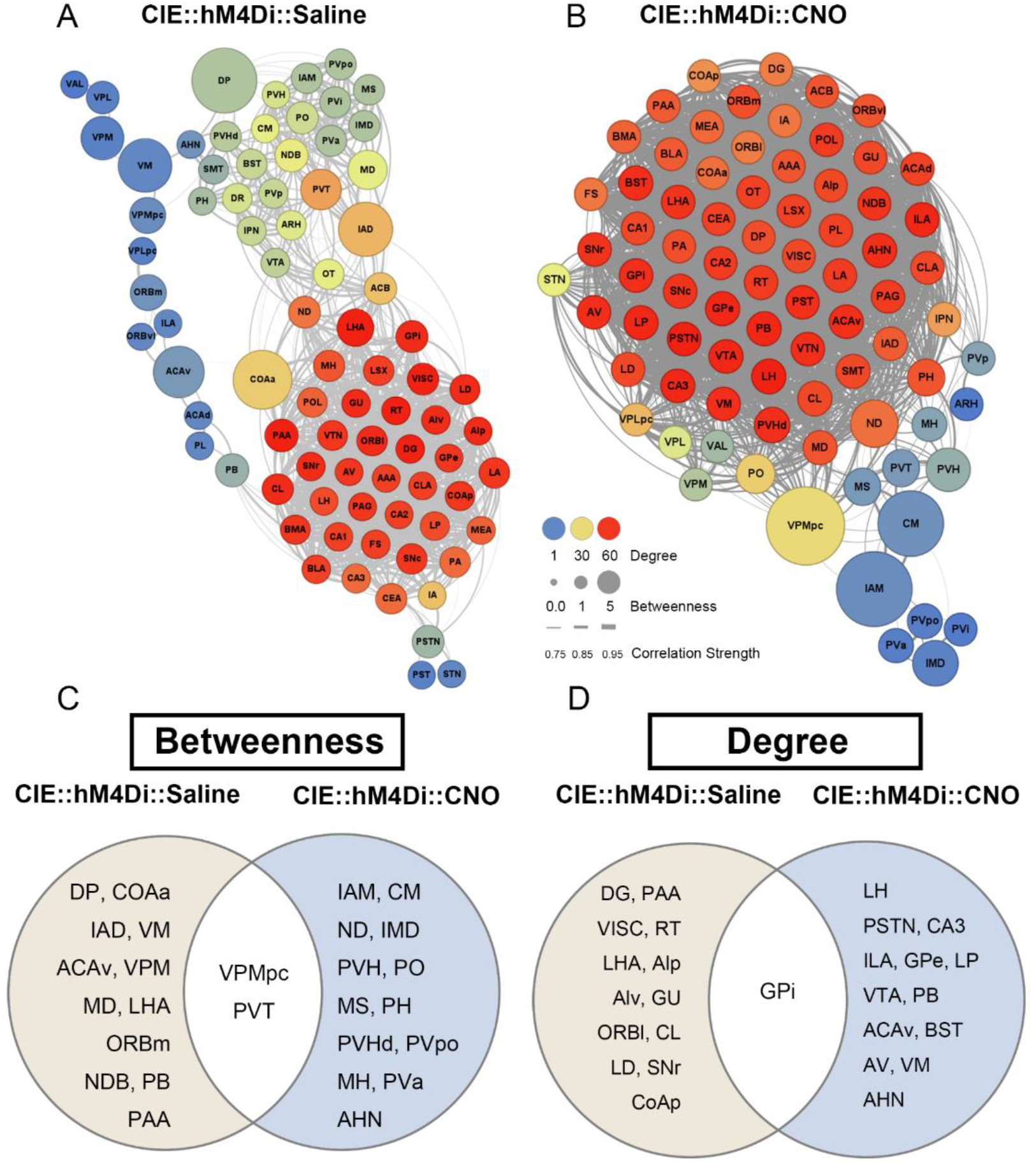
Neuronal networks of betweenness and degree are altered following cortical amygdala inhibition in alcohol-dependent mice. **(A-B)** Functional connectivity networks from CIE::hM4Di mice following saline or CNO. Node size indicates betweenness centrality; node color indicates degree. **(C-D)** Venn diagrams showing regions with significant changes in betweenness and degree centrality. Regional lists and centrality values are provided in Table 1-2. Full names of abbreviations are provided in Table S1.

Analysis of the top 15 hub regions ranked by betweenness centrality (Figure 5C; Table 1) showed that VPMpc and PVT (full names of abbreviations are provided in Table S1) consistently exhibited high centrality across both conditions, indicating that these two regions remained important network nodes regardless of whether the CoA was inhibited. Beyond these overlapping regions, distinct network patterns emerged between treatment conditions. In CIE::hM4Di::Saline group, key hub regions included DP, CoAa, IAD, VM, and ACAv. In contrast, following CNO-mediated inhibition of the CoA, thalamic regions such as IAM and CM, as well as paraventricular hypothalamic subregions (PVH, PVHd, and PVpo), became more prominent. Importantly, CoAa was a major hub during normal alcohol withdrawal but was no longer among the top-ranked regions after CoA inhibition. Instead, multiple thalamic nuclei gained prominence, suggesting that when CoA activity was suppressed, the thalamus regained a more central role in network connectivity during withdrawal. This observation is consistent with our earlier findings (Figure 7E–I) and supports the idea that suppressing CoA activity may restore thalamic engagement and thereby alleviate alcohol withdrawal-related network disruption.

**Table 1.** Brain regions with the highest betweenness centrality during alcohol withdrawal with or without CoA inhibition. Top 15 regions ranked by betweenness centrality in CIE mice expressing hM4Di-mCherry after saline or CNO treatment. Higher values indicate greater network influence. Regions unique to each condition are shown in black and shared regions in red. Full names of abbreviations are provided in Table S1.

| Betweenness |  |  |  |
| --- | --- | --- | --- |
| Saline |  | CNO |  |
| DP | 547.0323 | VPMpc | 209.9332 |
| COAa | 459.7109 | IAM | 196.202 |
| IAD | 391.9629 | CM | 152.1371 |
| VM | 379 | ND | 68.26976 |
| ACAv | 348.1949 | IMD | 59.2167 |
| VPM | 232 | PVH | 51.46266 |
| PVT | 194.2598 | PO | 27.9169 |
| MD | 170.2524 | MS | 20.71003 |
| LHA | 141.2812 | PH | 19.14083 |
| VPMpc | 129.1745 | PVT | 19.08118 |
| ORBm | 126.5992 | PVHd | 11.9119 |
| NDB | 86.71307 | PVpo | 8.291261 |
| PB | 84.57537 | MH | 7.949180 |
| PAA | 78.34854 | PVa | 7.790888 |
| VPL | 78 | AHN | 6.940991 |

Further analysis of the top 15 regions ranked by degree centrality also revealed additional network differences (Figure 5D, Table 2). Although GPi remained highly connected in both conditions, CoAp and PAA were hub regions during saline withdrawal but were no longer among the major hubs after CoA inhibition. Instead, LH, ILA, LP, bed nucleus of the stria terminalis (BST), VTA, and anterior hypothalamic nucleus (AHN) emerged as highly connected regions following CNO treatment. Notably, AHN ranked highly in both betweenness and degree centrality after CoA inhibition (Figures 5C-D), suggesting that AHN becomes a prominent network node when CoA activity is suppressed during alcohol withdrawal. Of note, the CoAp was a key region based on degree during the normal alcohol withdrawal state but was no longer prominent following CoA inhibition. The PAA, which was identified as a central region based on both betweenness and degree in saline-treated mice, was also absent from the top-ranked regions after inhibition. Overall, these findings suggest that chemogenetic inhibition of the CoA substantially alters the brain’s network architecture during alcohol withdrawal, promoting a shift in connectivity toward thalamic and isocortex regions.

**Table 2.**
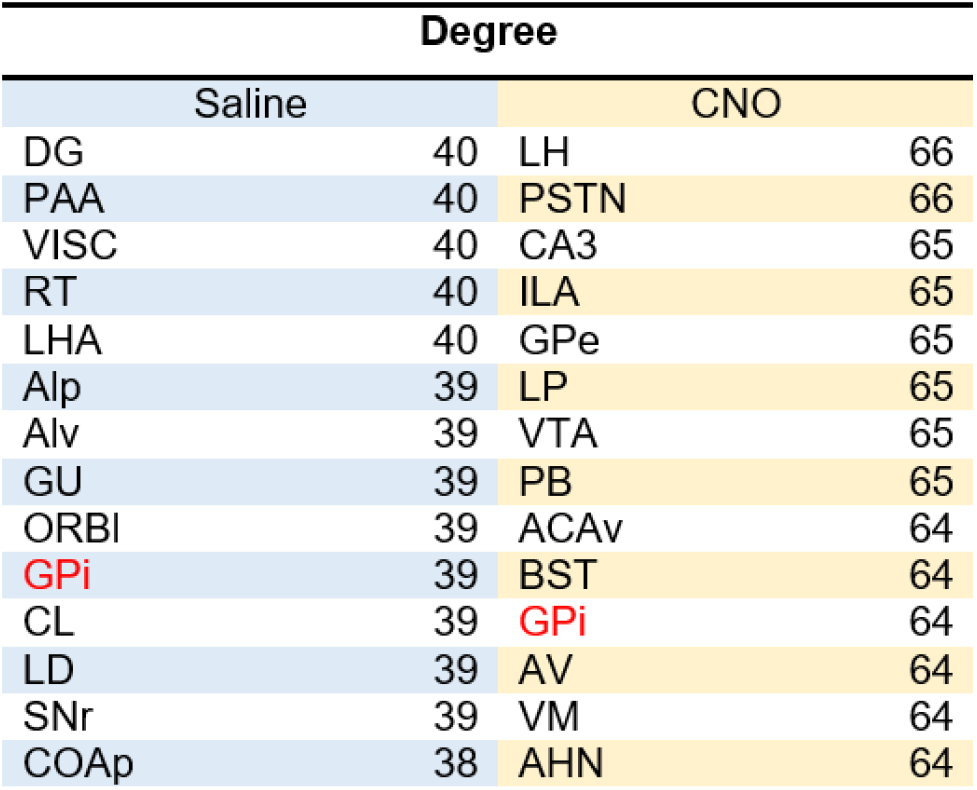
Brain regions with the highest degree centrality during alcohol withdrawal with or without CoA inhibition. Top 15 regions ranked by degree centrality in CIE mice expressing hM4Di-mCherry following saline or CNO treatment. Higher values indicate greater network connectivity. Regions unique to each condition are shown in black and shared regions in red. Full names of abbreviations are provided in Table S1.

## Discussion

In this study, electrophysiological recordings showed that CoA neurons from alcohol-dependent mice exhibited increased input resistance without changes in baseline firing rate, indicating altered intrinsic membrane properties during alcohol withdrawal. Acute ethanol selectively increased firing frequency and modified input resistance in neurons from alcohol-dependent mice, but not controls, suggesting that chronic alcohol exposure enhances CoA neuronal sensitivity to ethanol. Consistent with these cellular adaptations, we found that the CoA regulates alcohol-dependent drinking in female mice, extending previous male-focused findings (Roland et al., 2023). Although global inhibition of the CoA during withdrawal did not alleviate hyperalgesia or negative affect-like behaviors, it reduced overall network activity while strengthening functional connectivity. This shift restored disrupted thalamic communication and redistributed network hubs toward subcortical regions.

The increased excitability and ethanol sensitivity of CoA neurons are consistent with withdrawal-related adaptations reported in other alcohol-sensitive circuits. Similar changes in neuronal excitability have also been observed in other brain regions, suggesting that alcohol withdrawal may induce comparable neuroadaptations across multiple alcohol-sensitive circuits. Related withdrawal-associated adaptations have been reported in other extended amygdala structures, including enhanced excitatory signaling in corticotropin-releasing factor (CRF) neurons within the central amygdala (CeA) during withdrawal (Roberto et al., 2021), and increased intrinsic excitability in BST following withdrawal (Pati et al., 2020). In contrast, some neuronal populations, such as CRF neurons in the medial prefrontal cortex (mPFC), show reduced excitability during withdrawal (Patel et al., 2022), highlighting region-specific adaptations across alcohol-sensitive circuits. Overall, our findings support the idea that alcohol withdrawal promotes hyperexcitability in CoA neurons. This heightened responsiveness may alter circuit function and contribute to large-scale network reorganization that promotes maladaptive alcohol seeking and relapse-related behaviors.

Behaviorally, chemogenetic inhibition of the CoA reduced alcohol intake in dependent but not non-dependent mice. Our results therefore extend the role of the CoA in alcohol-dependent drinking across sexes. In contrast, CoA inhibition did not reduce withdrawal-related behaviors, including hyperalgesia, anxiety-like behavior, or depression-like behavior. These findings suggest that CoA activity primarily regulates alcohol consumption rather than broader behavioral symptoms of withdrawal. Alternatively, withdrawal-related behaviors may depend on projection-specific CoA circuits or interactions with other regions that were not targeted by the global chemogenetic inhibition used here.

At the network level, CoA inhibition produced widespread decreases in Fos expression during withdrawal, indicating that CoA hyperactivity contributes to elevated brain-wide activity in this state. Despite this reduction in overall neural activity, CoA inhibition increased co-activation across brain regions, suggesting improved coordination within the network. In particular, connectivity between the CoA and several isocortical regions (DP, GU, ILA, ORBL, ORBM, PAA, PL; full names of abbreviations are provided in Table S1) was enhanced, including multiple mPFC-associated areas (ILA, PL, and DP). mPFC, which plays a central role in executive functions such as cognition, memory formation, and decision-making (Euston et al., 2012; Grossmann, 2013; Wood and Grafman, 2003), and is also strongly implicated in alcohol-related behaviors (Heilig et al., 2017; Kroener et al., 2012). Since the mPFC projects to the CoA (Liu et al., 2024; Verwer et al., 1996), the enhanced mPFC–CoA connectivity may reflect restoration of top-down cortical control over alcohol consumption.

The thalamus was also strongly affected by CoA inhibition. During withdrawal, several thalamic nuclei showed negative co-activation with the rest of the brain, indicating disrupted thalamic integration. This observation is consistent with reports of thalamic hypoactivity in individuals with alcohol dependence and other substance use disorders (Huang et al., 2018; Salzwedel et al., 2016; Segobin et al., 2019; Tomasi et al., 2007; Zhang et al., 2016). Following CoA inhibition, these patterns shifted toward positive co-activity, suggesting improved thalamic coordination within the broader network. Network analysis further showed major hub reorganization: during normal withdrawal, cortical regions and the CoA itself emerged as prominent hubs, whereas CoA inhibition shifted hub organization toward multiple thalamic nuclei and increased the prominence of the AHN. Together, these findings suggest that excessive CoA activity during withdrawal disrupts thalamic communication and promotes cortical dominance, whereas suppressing CoA activity reorganizes brain-wide connectivity and enhances the contribution of thalamic circuits.

Together, these findings indicate that excessive CoA activity during withdrawal may disrupt thalamic communication, whereas suppressing CoA activity reorganizes brain-wide connectivity toward greater thalamic involvement. However, whether this reorganization restores connectivity toward a nondependent baseline remains unclear. In addition, because chemogenetic inhibition targeted the entire CoA, the specific neuronal populations or projection-defined circuits responsible for these effects could not be determined. Future studies using projection- or cell-type–specific approaches will be necessary to identify the precise CoA circuits that regulate alcohol consumption and large-scale network organization.

## Acknowledgments

This work was supported by National Institute of Health grants F.10025790.02.003 and F.10025790.02.001 by AK. The authors declare no conflicts of interest.

## Disclosures

The authors report no biomedical financial interests or potential conflicts of interest.

## Notes

### Competing Interest Statement

The authors have declared no competing interest.

