## Supplemental Figures for "Chemogenetic Inhibition of the Cortical Amygdala Reduces Alcohol Intake and Restores Thalamic Connectivity in Dependent Female Mice"

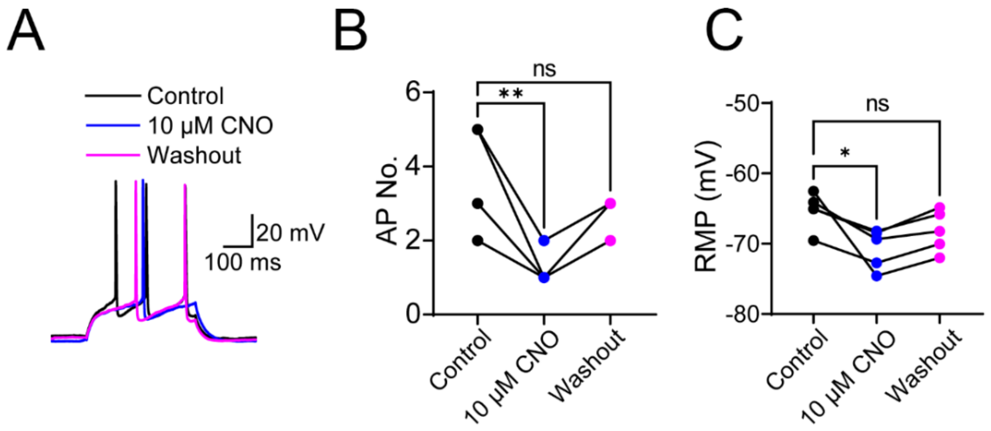


**Figure S1.** Validation of chemogenetic inhibition in hM4Di-expressing CoA neurons.
**(A)** Representative traces showing action potentials recorded under control conditions (black), during bath application of 10 μM CNO (blue), and after washout (purple).
**(B)** Quantification of the number of action potentials (AP No.) across individual neurons under control, CNO, and washout conditions. CNO significantly reduced action potential firing compared to control ($p < 0.01$), with firing returning toward baseline following washout (ns).
**(C)** Quantification of resting membrane potential (RMP) across conditions. CNO application resulted in a significant hyperpolarization compared to control ($P < 0.05$), which partially reversed after washout (ns). Data are shown as paired individual values with mean ± SEM. Statistical analysis was performed using repeated-measures ANOVA with Sidak’s multiple comparisons test.

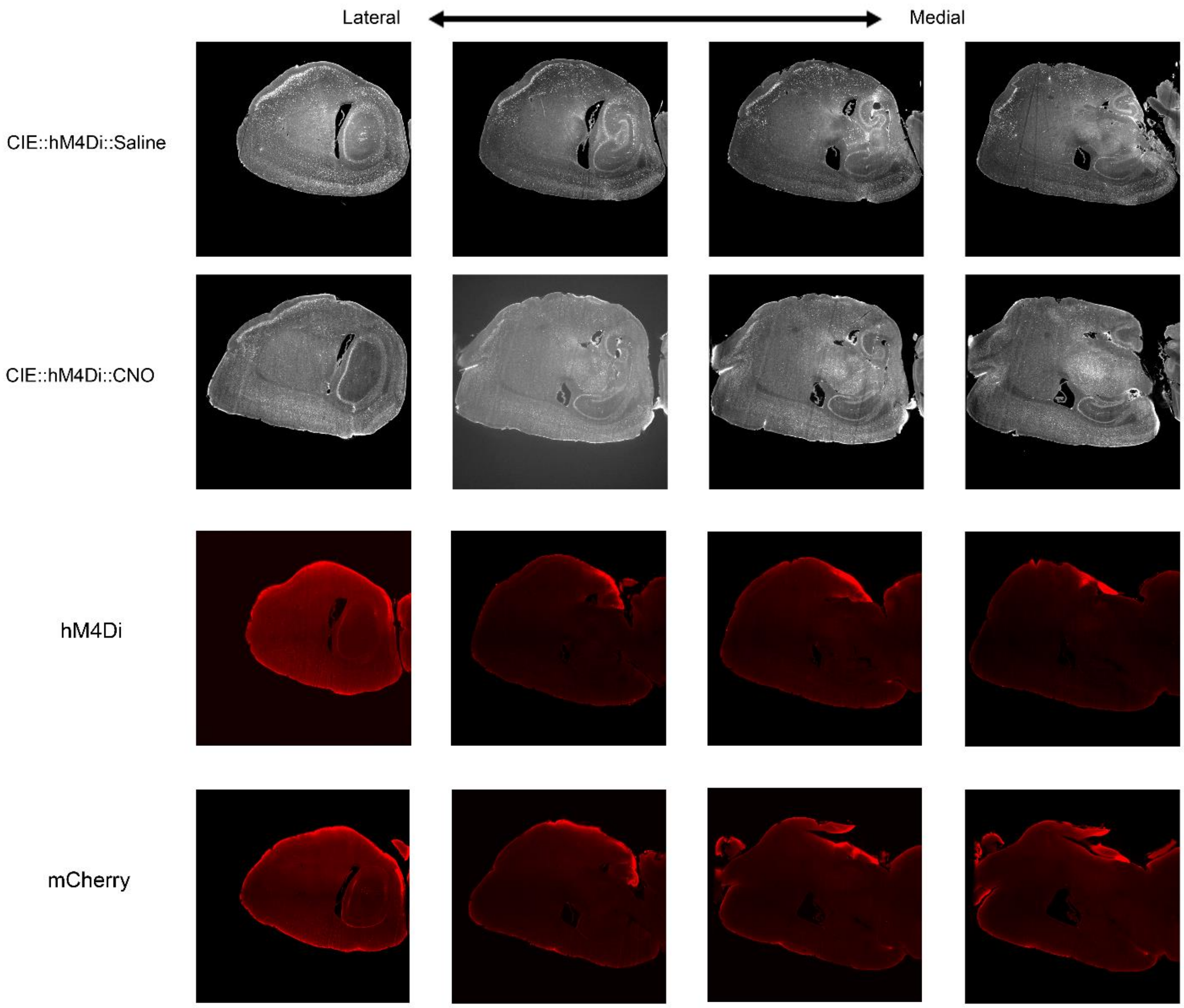


**Figure S2.** Representative images of Fos-labeled, iDISCO-cleared brain hemispheres in CIE::hM4Di mice treated with CNO or Saline.

Sagittal sections are shown from lateral to medial planes, with the top row depicting the CIE::hM4Di::CNO group and the bottom row depicting the CIE::hM4Di::Saline group. The bottom two rows show iDISCO-cleared fluorescent images of viral expression: hM4Di virus (third row) and sham control virus (mCherry, fourth row). These images highlight differences in c-Fos expression and neuronal activation patterns across the brain under conditions of CoA inhibition (CNO) compared to control (saline).

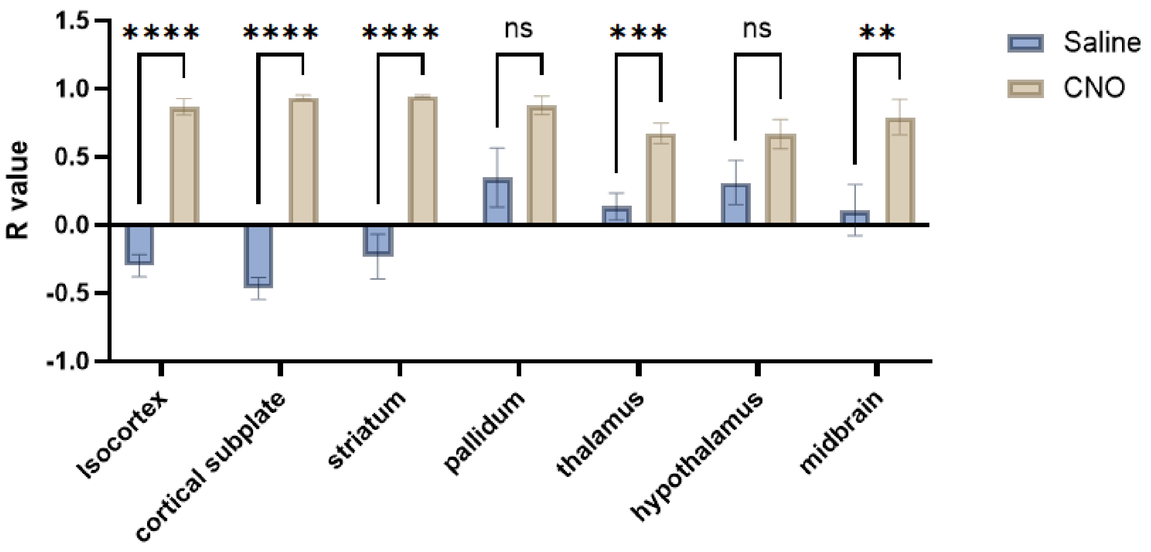


**Figure S3.** Chemogenetic inhibition of the Cortical Amygdala alters neuronal activity of AHN and functional connectivity associated with alcohol withdrawal
Bar graphs showing correlation coefficients (R values) of anterior hypothalamic nucleus (AHN) connectivity with major brain regions, including the isocortex, cortical subplate, striatum, pallidum, thalamus, hypothalamus, and midbrain, in CIE::hM4Di mice treated with saline or CNO. Chemogenetic inhibition of the CoA significantly increased functional connectivity between the AHN and these major regions. Statistical analysis was performed using two-way ANOVA followed by post hoc Sidak's comparison test ($P < 0.05$, $*P < 0.01$, $***P < 0.0001$).
